# Dual spindle formation in zygotes keeps parental genomes apart in early mammalian embryos

**DOI:** 10.1101/198275

**Authors:** Judith Reichmann, Bianca Nijmeijer, M. Julius Hossain, Manuel Eguren, Isabell Schneider, Antonio Z. Politi, Maria J. Roberti, Lars Hufnagel, Takashi Hiiragi, Jan Ellenberg

## Abstract

At the beginning of mammalian life the genetic material from each parent meets when the fertilized egg divides. It was previously thought that a single microtubule spindle is responsible to spatially combine the two genomes and then segregate them to create the two-cell embryo. Utilizing light-sheet microscopy, we showed that two bipolar spindles form in the zygote, that independently congress the maternal and paternal genomes. These two spindles aligned their poles prior to anaphase but kept the parental genomes apart during the first cleavage. This spindle assembly mechanism provides a rationale for erroneous divisions into more than two blastomeric nuclei observed in mammalian zygotes and reveals the mechanism behind the observation that parental genomes occupy separate nuclear compartments in the two-cell embryo.

**One Sentence Summary:** After fertilization, two spindles form around pro-nuclei in mammalian zygotes and keep the parental genomes apart during the first division.

---

After fertilization, the haploid genomes of egg and sperm come together to form the genome of a new diploid organism, a moment that is of fundamental biological importance. In mammals, parental chromosomes meet for the first time upon entry into the first zygotic mitosis after nuclear envelope break down (NEBD). So far, it was assumed that similar to oocytes a single bipolar microtubule system would self-assemble around both parental genomes also in zygotes (*1–6*). Due to the extreme light-sensitivity of the mammalian embryo the details of the dynamic process of zygotic spindle assembly, however, remained unclear.

To examine how parental genomes join for the first time, we imaged live embryos in which the maternal and paternal centromeres were differentially labelled (*7*) using our recently developed inverted light-sheet microscope, which allows fast 3D imaging of embryonic development due to its low phototoxicity (*8*). This revealed that the two genomes remain spatially separate throughout the first mitosis (Fig. S1, Movie S1). To understand why the genomes are not mixed, we next imaged spindle assembly using fluorescently labelled microtubule organizing centers (MTOCs) and spindle microtubules (Fig. 1A, Movie S2). We found that newly nucleated microtubules self-organized into two separate bipolar spindles after NEBD attracting a subset of the cytoplasmic MTOCs that had accumulated around each pronucleus to their poles (Fig. 1A, B). Subsequently, the two spindles aligned and came into close apposition to form a compound barrel-shaped system. This structure typically had two clusters of MTOCs at at least one of its poles, suggesting that the two spindles were aligned closely but not completely merged (Fig. 1A, B, Movie S2). To probe this further, we performed 3D immunofluorescence analysis of zygotes visualizing endogenous spindle poles, microtubules, kinetochores and DNA. This showed that in early and mid prometaphase, two separate bipolar spindles are formed in *in vivo* developed zygotes (Fig. 1C, D). Given the delayed and partial association of MTOCs with the microtubule mass, we hypothesized that dual spindle formation might be driven by self-assembly of microtubules nucleated by chromosomes. To test this, we assayed microtubule regrowth after washing out the microtubule depolymerizing drug nocodazole. Indeed, a large proportion of microtubules was nucleated on chromosomes with striking association to kinetochores (Fig. S2, while MOTCs became associated with microtubules only later. This observation prompted us to investigate the organization of K-fibers in zygotic prometaphase by high-resolution immunofluorescence of zygotes fixed after brief cold treatment to highlight stable microtubules (Fig. S3). This analysis showed that two bipolar arrays of K-fibers start assembling in early prometaphase, are stably organized in mid prometaphase, and clearly recognizable by their slightly offset centers and split poles even after parallelization in metaphase.

**Figure 1.**
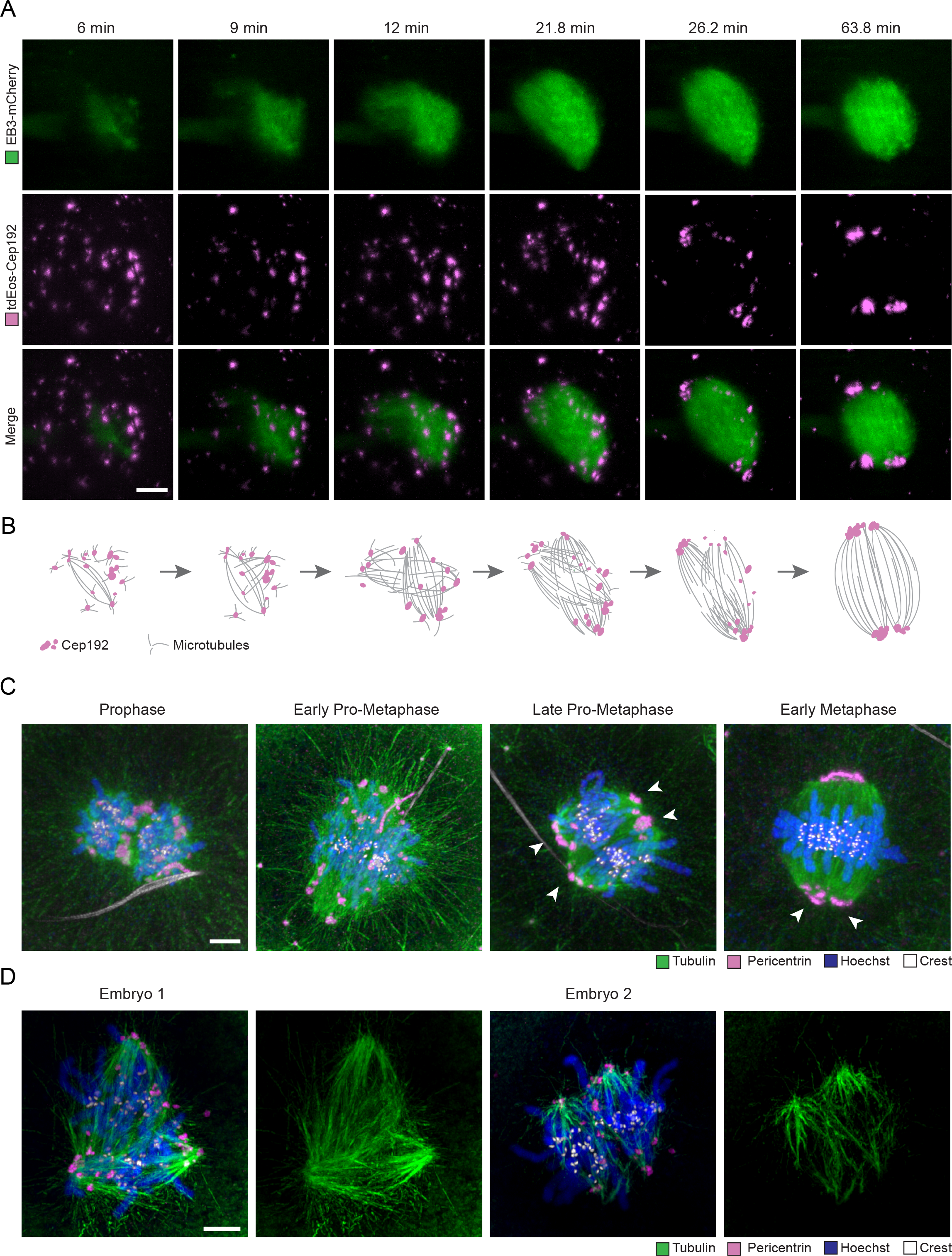
Individual bi-polar spindle formation around each pronucleus. (A) Time-lapse-imaging of *Mus Musculus* x *Mus Musculus* (MMU x MMU) zygotes expressing EB3-mCherry (marker for microtubules; green) and tdEos-Cep192 (marker for MTOCs; magenta). Scale bar, 10 µm. In 11 out of 13 zygotes both or at least one of the dual spindle poles remained clearly split after the spindles had parallelized. (B) Schematic of progression of dual spindle formation. Microtubules in grey, MTOCs in magenta. (C) Immunofluorescence staining of MMU x MMU zygotes fixed at consecutive stages of development. Shown are z-projected images of confocal sections of zygotes at prophase; early pro-metaphase, late pro-metaphase, early metaphase; late metaphase and anaphase. (D) Immunofluorescence staining of cold treated MMU x MMU prometaphase zygotes. Shown are z-projected images of confocal sections. Microtubules (Tubulin; green), MTOCs (Pericentrin; magenta), kinetochores (Crest; white), and DNA (Hoechst; blue) are shown. Scale bars, 5 µm (A, C, D). White arrows indicate poles (A, C).

To characterize the kinetics of zygotic spindle assembly in live embryos, we next imaged maternal and paternal centromeres in relation to growing spindle microtubule tips. This allowed us to define three phases of zygotic spindle assembly (Fig. 2A). A transient first phase (~3 min; 10.3 ± 3.5 min to 13. 4 ±4 min after NEBD), characterized by the clustering of growing microtubules around the two pronuclei; followed by phase 2 (~16 min; 14.5± 4 min to 30.7 ± 6.5 min after NEBD), where individual bipolar spindles assembled around each parental genome; and subsequently phase 3 (~83 min; 46.7±17 min to 129.2 ± 16.5 min after NEBD), when the two spindles align and combine into a compound barrel shaped structure.

**Figure 2.**
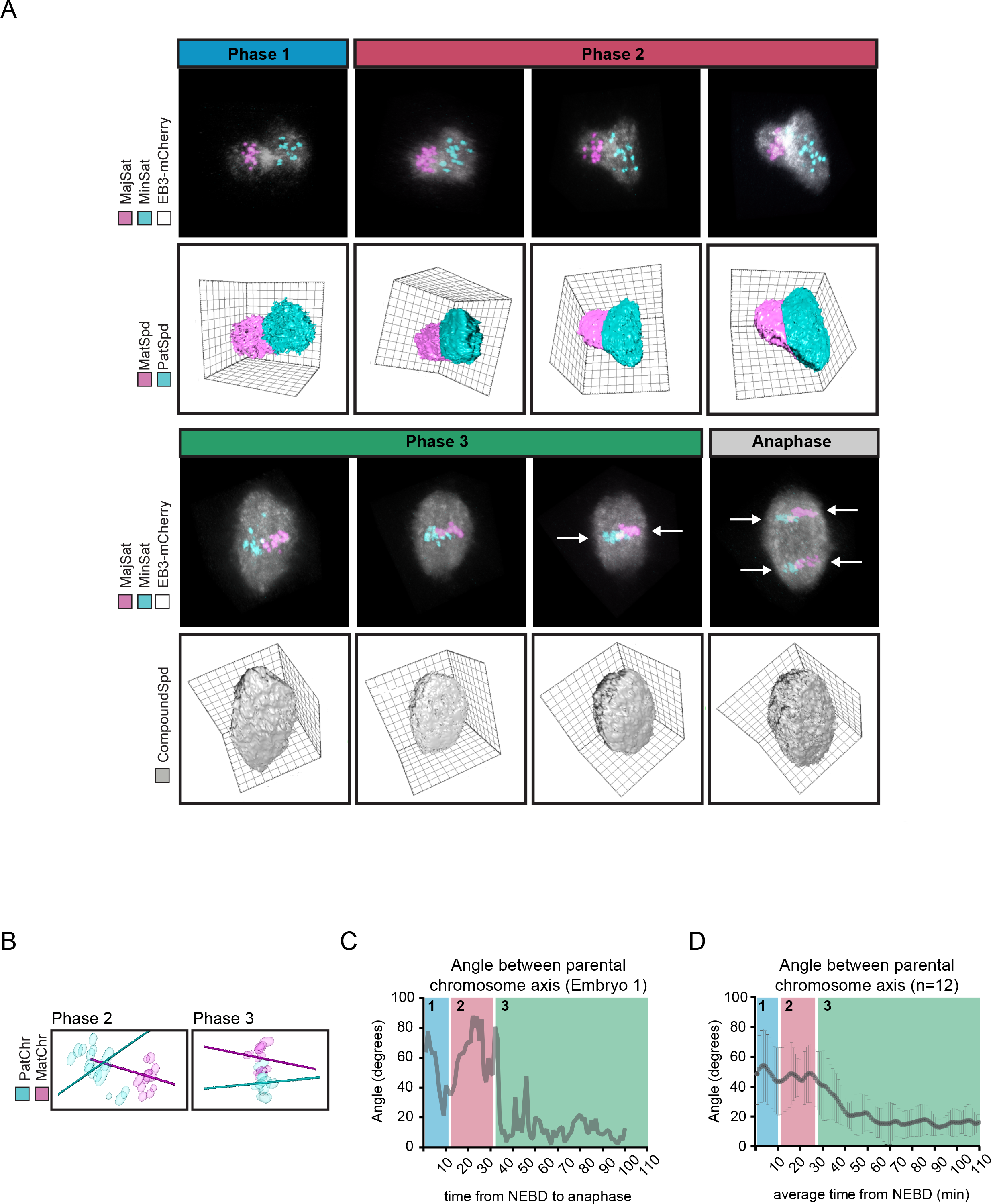
Spindle assembly and chromosome dynamics in the zygote. (A) Time-lapse-imaging of *Mus Musculus* x *Mus Spretus* (MMU x MSP) zygotes expressing EB3-mCherry and fluorescent TALEs to label maternal (MajSat) and paternal (MinSat) chromosomes. Phase 1: Microtubule ball formation around pronuclei. Phase 2: Bi-polarization of maternal and paternal spindle. Phase 3: Formation of single barrel-shaped spindle. Upper rows, 3D-rotated images of the whole spindle volume. Maternal (MajSat; magenta) and paternal (MinSat; cyan) chromosomes, and microtubules (EB3-mCherry; white). Lower rows, segmentation of maternal (MatSpd; magenta) and paternal (PatSpd; cyan) spindles in Phase 1 and Phase 2 and single bipolar spindle in Phase 3 (CompoundSpd; grey). Offset between maternal and paternal chromosomes at metaphase and anaphase is indicated with white arrows. (B) Schematic of measurements on maternal (MatChr) and paternal chromatin masses (PatChr) in phase 2 and 3. (C,D) Angles between maternal and paternal chromosome axis over time for a single embryo (C) and averaged for 12 embryos (D; mean ± SD) are shown. Phase 1, blue; Phase 2, red; Phase 3, green.

To test if the two zygotic spindles are functionally independent, we measured the timing and direction of maternal and paternal chromosome congression (Fig S.4A, B and Fig. 2A, B; for details see methods). Congression started in prometaphase (phase 2), while the spindles were clearly separated (Fig. S4A, B). Parental chromosome congression was not correlated in time until shortly before anaphase, suggesting that they are moved by different microtubule systems (Fig. S4C). Furthermore, the parental genomes were congressed in different directions along separate spindle axes, as evidenced by the large difference between the angles of the two forming metaphase plates, which became parallel only later during dual spindle alignment in phase 3 (Fig. 2B-D; Fig. S4D, E). Tracking of growing microtubule tips showed two major directions of microtubule flow during phase 2 and one main direction during phase 3 as indicated by the corresponding kymograph profiles (Fig. S5, Movies S3-8). The independent congression frequently led to an offset in the final bi-oriented position of the paternal and maternal metaphase plates at the end of phase 3 prior to and during segregation (Fig. 2A, arrows). Together, this data suggests that each of the two spindles around the parental genomes functions independently for chromosome congression, and that they may even function uncoupled from each other in chromosome segregation.

It has been observed in in-vitro-fertilization (IVF) clinics, that the zygotic division is error prone and often leads to embryos with blastomeres containing two nuclei (*9–13*). Based on our observations, we hypothesized that failure to align the two parental spindles before anaphase could explain this enigmatic phenotype. In order to test this, we increased the distance between the two pronuclei by transient treatment with nocodazole, which led to a larger gap between the two self-assembling spindles (Fig. 3, Movies S9-11). Indeed, such embryos frequently failed to fully align the parental spindles at one or both poles. Strikingly, this did not delay anaphase but resulted in chromosome segregation by two spindles into different directions, leading to two cell embryos with one or two bi-nucleated blastomeres (Fig. 3, Movies S10-11). By contrast, embryos that did align the two spindles parallel to each other before anaphase cleaved into two blastomeres with single nuclei as expected (Movie S9). Thus, failure to align the two zygotic spindles gives rise to multinucleated two-cell embryos, phenocopying frequently observed errors in human embryonic development in IVF clinics.

**Figure 3:**
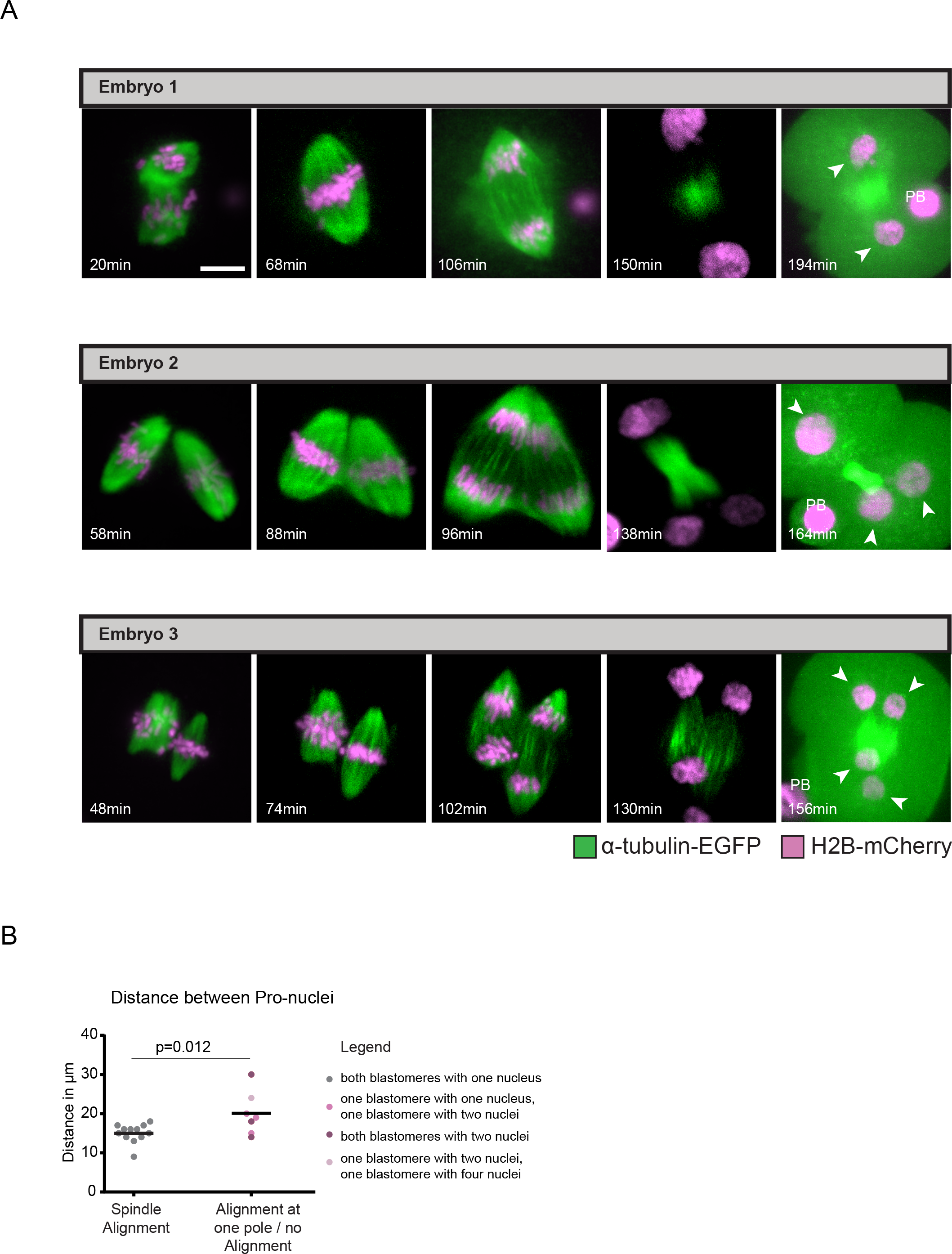
Proximity dependency of bipolar spindle fusion. (A) Time-lapse imaging of MMU x MMU zygotes expressing H2B-mCherry (chromatin; magenta) and αTubulin-EGFP (microtubules; green). Spindle morphology in pro-metaphase and anaphase in 3 representative zygotes treated with Nocodazole for > 10 hours. *z*-projected images of pro-metaphase, metaphase, anaphase, telophase and post-mitosis are shown. Arrowheads and PB indicate nuclei and polar body, respectively. Scale bar, 10 µm. In absence of NEBD as timing reference, anaphase onset was set at 90 minutes (average time from NEBD to anaphase in MMU x MMU zygotes) and the other times calculated accordingly. (B) Initial distance of pro-nuclei. Statistics, students’ t-test.

Dual spindle assembly in the mammalian zygote would also offer a mechanistic explanation for the long-standing observation that the parental genomes occupy separate compartments inside the nuclei of two- and four-cell embryo blastomeres (*14, 15*). If dual spindle assembly around two pronuclei was responsible for genome compartmentalization, parental genomes should mix in subsequent divisions, where only one nucleus is present per cell. Imaging of the metaphase plate of live hybrid mouse embryos from the zygote to the eight-cell stage showed that parental genomes were separated in zygotes, but became rapidly mixed in the subsequent developmental stages as predicted (Fig. S6A-D). This loss of genome compartmentalization was also seen in *in vivo* developed isogenic embryos (Fig. S6E-I). Thus parental genomes are kept separate by two spindles only during the first mitosis but then mix during subsequent divisions driven by a single common spindle.

If dual spindle assembly is the mechanism for parental genome compartmentalization (Fig. S7A), formation of a single spindle around both genomes in the zygote should mix them already in the first division. To test this prediction, we redirected spindle assembly with two small molecule inhibitors of microtubule polymerization (Nocodazole) and the motor protein Eg5 (Monastrol). Transient treatment with Monastrol collected both genomes in a single microtubule aster and subsequent depolymerization of microtubules with Nocodazole followed by regrowth after washout then resulted in one bipolar spindle around both genomes (Fig. S7B and S8, from here on referred to as MoNoc treated zygotes). Such MoNoc treated embryos captured and congressed chromosomes within a single spindle and showed a high degree of parental genome mixing in the first mitotic metaphase (Fig. 4). This was significantly different from untreated or control zygotes in which the order of drug treatments is reversed (NocMo treated zygotes), which maintained dual spindle formation and genome separation (Fig. 4; Fig. S7C and S8). Thus, dual spindle formation in the zygote is required for parental genome separation in mammals. Having this experimental method to induce mixing of the parental genomes in hand, furthermore allowed us to demonstrate that genome separation is not required for the epigenetic asymmetry, and its resolution (*16–19*) between parental genomes as proposed previously (Fig. S9 and S10) (*15, 20–22*), which is thus a chromosome intrinsic property.

**Figure 4.**
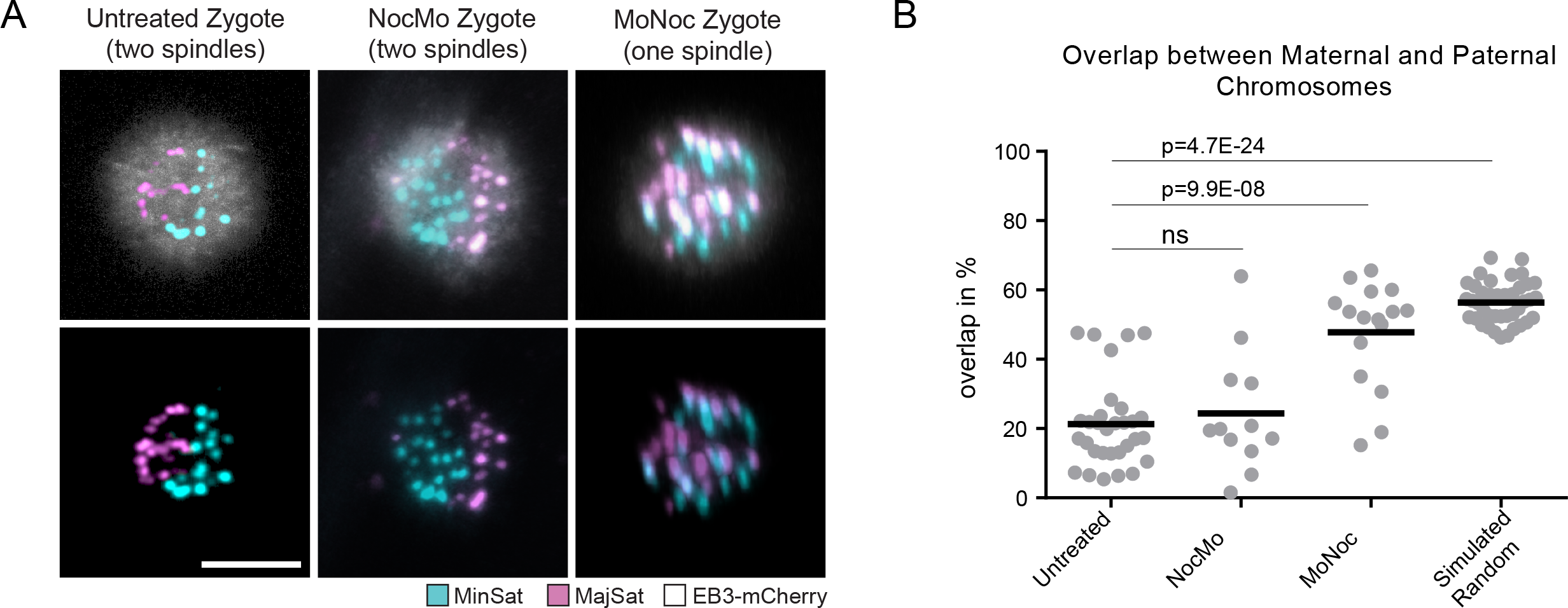
Distribution of parental centromeres in control, NocMo and MoNoc treated zygotes. (A) Differential labelling of maternal (MajSat; magenta) and paternal (MinSat; cyan) centromeres through distinction of SNPs by fluorescent TALEs. Mitotic spindle is labelled with EB3-mCherry (white). Representative *z*-projected images of parental chromosome distribution in untreated, MoNoc and NocMo MMU x MSP zygotes. Scale bar, 10 µm. (B) Degree of overlap between 3D convex hulls and parental chromosomes for untreated (*n* = 31), MoNoc (*n* = 16), NocMo (*n* = 12) zygotes and embryos with *in silico* randomized distribution (*n* = 40) (see Figure S1 and methods for details). Statistics, students’ t-test.

In summary, we showed here that two spindles form around pronuclei in mammalian zygotes which individually collect the parental genomes and then position them next to each other prior to the first anaphase. Our data explain how parental genome separation is achieved in mammalian embryos. To date, the formation of physically distinct mitotic spindles around the two pronuclei has been thought to be specific to certain arthropod species (*23, 24*). Our finding that this occurs also in mammals, suggests that two zygotic spindles might be characteristic for many species that maintain separate pronuclei after fertilization. We demonstrated that failure to align the two spindles produces errors in the zygotic division that closely resemble clinical phenotypes of human embryos in IVF procedures, suggesting that a similar mechanism of dual zygotic spindle assembly also occurs in human. This view is supported by the spatial separation of parental chromosomes reported in human zygotes (*25*), divisions of human zygotes into multinucleated blastomeres and by reports that identified a mix of paternal, maternal and diploid cells in eight-cell cattle embryos. A dual zygotic spindle would provide a mechanistic basis for this parental genome segregation (*9–13, 26, 27*). These severe and relatively frequent zygotic division errors in human and agriculturally used mammals thus find their likely mechanistic explanation in a failure of the close alignment of the two zygotic spindles prior to anaphase. Beyond the biological insight, if a similar mechanism of microtubule-driven parental genome separation indeed occurs in human zygotes would be important to understand from an ethical and legal perspective as “pronuclear fusion”, a process that strictly speaking does not occur in mouse zygotes, is used to define the beginning of embryonic life as protected by law in several countries (e.g. Germany, § *8 Abs*. *1* Embryonenschutzgesetz).

## Acknowledgments

We thank Nathalie Daigle for cloning of the EB3-mCherry plasmid, Pierre Neveu and Melina Schuh for providing tdiRFP670 and tdEos-Cep192, respectively, and Niels Galjart for providing full length *Homo sapiens* EB3 cDNA (NM_001303050.1). We thank Petr Strnad for development and assistance of the inverted light-sheet microscope, and Arina Rybina for assistance with the Zeiss LSM 880 Airy microscope. We thank EMBL’s laboratory of animal resources for support and the EMBL Advanced Light Microscopy Facility is acknowledged for support in image acquisition and analysis. We thank Arivis for support in image analysis. We thank James Reddington and Stephanie Alexander for critical reading of the manuscript.

## Author Contributions

J.E. and J.R. conceived the project and designed the experiments. J.R., B.N., M.E. and I.S. performed the experiments. M.J.R. supported the mouse EDU experiments. J.R., J.H. and A.P. analyzed the data. T.H. and L.H. contributed to conception and design of the work. J.E. and J.R. wrote the manuscript. All authors contributed to the interpretation of the data and read and approved the final manuscript.

## Data materials availability

All image data used in this manuscript are available from XXX. Scripts and codes are available from XXX.

## Supplementary Materials

Materials and Methods

Figs. S1 to S10

Table S1

References 28 - 34

Movies S1 to S9

## Materials and Methods

### Mouse strains and embryo culture

Mouse embryos were collected from superovulated 8- to 24-week-old female mice according to the guidelines of EMBL Laboratory Animal Resources and cultured in 30-µl drops of G1 (Vitrolife) covered by mineral oil (Ovoil, Vitrolife). Embryos used for immunofluorescence were isolated from C57BL/6J x C3H/He F1 females, or EGFP-Tuba C57BL/6J x C3H/He F1 females, mated with C57BL/6J x C3H/He F1 males and fixed at different stages of zygotic mitosis. Embryos used for imaging of parental chromosomes were isolated from C57BL/6J x C3H/He F1 or H2BmCherry C57BL/6J x C3H/He F1 females mated with *Mus Spretus* males (*Mus musculus* (MMU) and *Mus spretus* (MSP) hybrid embryo). Culture during imaging was performed as described (*8*) with minor modifications. In brief, embryos were imaged in G1 medium covered with mineral oil with 5% CO_2_ and 5% O_2_ atmosphere. To achieve mixing of chromosomes embryos were cultured with 0.1 mM Monastrol for 5 hours followed by 0.01 mM Nocodazole for 1 hour. Subsequently, embryos were imaged or allowed to develop until the two cell stage and then synchronized with 0.01 mM Nocodazole or 0.1 mM Monastrol and then fixed. For controls the order of drug treatment was reversed in addition to a no drug treatment control. To assay microtubule regrowth zygotes were cultured in presence of 0.01 mM Nocodazole until nuclear envelope breakdown (5 to 7 hours). Nocodazole was then washed out, embryos transferred back into culture medium and fixed in 1-min-intervals as described below.

For EdU labelling of paternal chromatin male from C57BL/6J x C3H/He mice were supplied continuously with drinking water containing 0.5 mg/ml EdU. The EdU-containing water was changed twice per week. EdU is incorporated in place of thymidine into the replicating DNA of mitotically dividing somatic and pre-meiotic cells. The cycle time to produce mature spermatozoa from EdU-labelled pre-meiotic cells in mice is ∼35 days. After continuous EdU feeding for at least 6 weeks, EdU-treated males were mated with superovulated B6C3F1 females. Embryos were isolated 20 hours after HCG injection and cultured in G1 medium (Vitrolife) up to the 8-cell stage.

Embryos used for live imaging of microtubule tips were isolated from C57BL/6J x C3H/He F1 females mated with C57BL/6J x C3H/He F1 males.

### Expression Constructs and mRNA Synthesis

Constructs used for mRNA synthesis were previously described: TALE-mClover (pTALYM3B15 Addgene plasmid 47878) (*7*), EB3-mEGFP (*4*), tdEos-Cep192 (*28*) (a kind gift from Melina Schuh). To generate EB3-mCherry full length Homo sapiens EB3 cDNA (NM_001303050.1, a generous gift from Niels Galjart) was tagged at the C-terminus with a tandem mCherry and cloned into the vector pGEMHE for mRNA production. To generate TALE-tdiRFP670, mRuby from pTALYM4SpiMi-01 (*7*) (Addgene plasmid 47879) was replaced with tdiRFP670 (Addgene plasmid 45466, the tandem construct was a kind gift from Pierre Neveu). After linearization of the template with PacI, capped mRNA was synthesized using T7 polymerase (mMessage mMachine Ultra Kit, following manufacturer’s instructions, Ambion) and dissolved in 11 µl water. mRNA concentrations were determined using a NanoDrop (Thermo Fisher Scientific).

### Immunofluorescence

For imaging of the mitotic spindle and to assay microtubule regrowth embryos were fixed and extracted as described (*5*). To visualize organization of K-fibers in zygotic prometaphase embryos were incubated in ice-cold PBS for 3 minutes prior to fixation. Embryos were blocked in 5% normal goat serum, 3% BSA in PBST (0.1% Trition X-100) and then incubated overnight in blocking solution at 4°C at the following antibody dilutions:1:500 mouse anti-tubulin (Sigma T6199) to visualise microtubules, 1:500 rabbit anti-pericentrin (Abcam ab4448) for staining of MTOCs, 1:100 human anti-Crest (Europe Bioproducts CS1058) to stain centromeres. Embryos were washed 3 x 5 minutes with 0.3% BSA in PBST then incubated with anti-mouse Alexa 488, anti-rabbit Alexa 546, anti-human Alexa 647 all 1:500 in 5% normal goat serum, 3% BSA in PBST (all (Thermo Fisher Scientific A11029, A11035, A21445 respectively) and 5ug/ml Hoechst33342 (Sigma) for 1 hour at room temperature. Embryos were washed with 0.3% BSA in PBST for 3 x 5 minutes before imaging.

To visualise EdU-substituted DNA embryos were fixed with 1%PFA for 30 minutes and then washed three times in PBS. For staining of EdU labelled DNA embryos were treated with Click-iT^®^ Labelling Technologies (Thermo Fisher Scientific C10340) according to the manufacturer’s instructions. Paternal chromatin was labelled with Alexa Fluor 647 and the whole nucleus counter stained with 100 nM Sytox Green (Molecular probes S7020).

For imaging of 5mC and 5hmC embryos were fixed for 20 minutes with 4% PFA in PBS. Embryos were washed 3 x in 1% BSA in PBS then extracted overnight in 1% BSA in PBS containing 0.5% Trition X-100. Embryos were incubated for 1 hour at 37 °C in the presence of 10 µg/ml RNAse A (Sigma). Chromosomes were denatured by incubating the embryos in 4N HCL for 30 minutes at 37°C followed by neutralisation with 100mM Tris buffer (pH8) at room temperature. Embryos were blocked with 3% BSA and 5% normal goat serum in PBST and then incubated at 4°C overnight with 1:3000 mouse anti 5-methylcytosine (Diagenode C152000081) and 1:3000 rabbit anti 5-hydroxymethylcytosine (RevMab Biosciences 31-1111-00). Embryos were washed with 3% BSA in PBST for 3x 5 minutes then incubated with 1:1500 anti-rabbit Alexa 647 (Thermo Fisher Scientific A21245), 1:1500 anti-mouse Alexa 546 (Thermo Fisher Scientific A11030) and 100nM Yoyo-1 (Thermo Fisher Scientific Y3601) for 1 hour at room temperature. Embryos were then washed 3 x for 5 minutes with 3% BSA in PBST and imaged.

For immunofluorescence of H3K9me3 and Ring1B embryos were fixed as described (*5*). Embryos were incubated overnight at 4 °C at the following primary antibody dilutions: 1:2000 rabbit anti Ring1B (Abcam 101273) or 1:200 rabbit anti-H3K9me3 (Abcam 8898) in 5% normal goat serum, 3% BSA in PBST. Embryos were washed with 0.3% BSA in PBST for 3 x 5 minutes then incubated with 1:1000 anti-rabbitAlexa 647 and 5ug/ml Hoechst (Sigma) for 1 hour at room temperature. Embryos were washed with 0.3% BSA in PBST for 3 x 5 minutes before imaging.

### Micromanipulation

Embryos were injected based on methods described previously (*4*). The injected volumes ranged between 10–15 pl (3%–5% of the embryo volume) of 0.125 – 0.3 µg/µl mRNA. mRNA-injected embryos were incubated at 37°C for 4–6 hr in G1 medium as described above to allow recombinant protein expression. For labelling of MTOCs and microtubule tips MMU zygotes were injected with mRNA encoding tdEos-Cep192 and EB3-mCherry. For differential labelling of maternal and paternal centromeres MMU x MSP embryos were injected with mRNA encoding fluorescent proteins fused to TALEs specific to the different centromeric satellite repeats as described previously (*7*).

### Embryo Imaging

Time-lapse image acquisitions were performed using a previously described in-house-built inverted light-sheet microscope (*8*) with the following modifications: (i) Image acquisition was performed with an Orca Flash 4 V2 camera from Hamamatsu Photonics, Japan. (ii) A ZYNQ-based cRIO-9064 embedded controller from National Instruments was used for real-time control of all microscope components. (iii) A rotating 6 mm thick and 25 mm diameter glass plate (Thorlabs) was inserted in the illumination path between the objective lens and the tube lens to translate the beam in the back focal plane of the illumination objective lens. For imaging MTOCs and microtubules stacks of 101 images with 520 nm between planes were acquired simultaneously for mCherry and EGFP signals at 45 sec time intervals. For imaging of growing microtubule tips 5 images with 500 nm between planes were acquired simultaneously for mCherry and EGFP at 800 ms intervals with an exposure time of 100 ms. Fixed EdU treated embryos were imaged by acquiring 101 images with 520 nm between planes consecutively for Sytox Green and Alexa 647 signals.

Fixed embryos stained for spindle components and epigenetic marks were imaged on a SP8 Leica confocal microscope equipped with a 63× C-Apochromat 1.2 NA water immersion objective lens. Images of embryos stained for spindle components or epigenetic marks were acquired at 90 nm in *xy* and 360 nm in Z. Images of embryos assayed for microtubule regrowth were acquired at 80 nm in *xy* and 320 nm in *z*.

Embryos cold treated prior to fixation, to assess organization of K-fibers in zygotic prometaphase, were imaged on a Zeiss LSM 880 confocal microscope with Airyscan equipped with a 40× C-Apochromat 1.2 NA water immersion objective lens. Images were acquired at 80 nm in *xy* and 220 nm in *z*.

### Image processing and analysis

Images of embryos stained and fixed for spindle markers were deconvolved using the Huygens remote manager (Scientific Volume Imaging), and maximum intensity projected in Arivis (arivis Vision4D). Images of pro-metaphase zygotes acquired on the Zeiss LSM 880 were subjected to 2D Airyscan processing and rotated in the Arivis 3D Viewer (arivis Vision4D) for presentation purposes.

Time-lapse images were processed for extraction of raw camera data as described (*8*). Time-lapse movies were generated as described (*8*) or exported from Arivis. Phases of zygotic mitosis were scored manually according to spindle morphology and presence of two bi-polar or single barrel shaped spindle. Spindles were segmented using an Arivis inbuilt intensity threshold filter. Chromosomes were segmented using an in house developed MATLAB segmentation pipeline based on intensity threshold and connected component analysis. Shape and direction of chromosomes are represented using Eigen value and Eigen vector of the segmented chromosomes in order to measure chromosome congression and the angle between parental chromosomes.

#### Segmentation and analysis of EdU signal

The nucleus was segmented from the DNA signal and used as a mask to segment the EdU signal. The center of nuclear mass was detected and a plane was fitted through the centroid to achieve maximum separation of the EdU signal in the two resulting hemispheres. Then, the volume of the smallest EdU signal was divided by the volume of the largest EdU signal.

#### Segmentation and analysis of maternal and paternal genomes

The chromosome mass was segmented from the DNA channel and used as mask to crop maternal and paternal chromosome signals. These signals were then segmented using intensity threshold without considering the regions masked out. Segmented maternal and paternal chromosomes were bounded by two separate 3D convex hulls. The mixing between the two was computed by the volume of overlapping voxels between two convex hulls divided by the volume of the union of the two convex hulls.

#### Simulation of random distribution of paternal chromatin signal at interphase

Three spheres were generated representing chromosome masses of 2-cell, 4-cell and 8-cell embryos. The average volume of cells in each stage was calculated from the original data in order to select the radius of each sphere to achieve comparable relative volumes of chromosome masses in different stages. 40, 20 and 10 small spheres representing paternal chromatin signal were generated with the chromosome masses of 2-cell, 4-cell and 8-cell embryos, respectively.

#### Simulation of random distribution of maternal and paternal centromere signals at metaphase

A model for chromosome mass was generated by combining the shape information of several individual metaphase chromosome masses. 40 spheres with the same radius were generated in random location within the model chromosome mass of which 50% represent maternal and 50% paternal chromosomes. A minimum distance between the centroids of any two spheres was enforced in order to avoid generating multiple spheres in the same location.

#### Correlation of maternal and paternal chromosome congression (15-frame window)

To segment chromosomes, original images were filtered first using a 2D Gaussian filter of size 3 and SD=2. Filtered images were interpolated along Z to obtain isotropic pixel size. Pixels at a distance from the center of mass >13 µm were discarded to reduce the effect of noisy blobs to be detected as chromosomes. Each slice of the interpolated stack was binarized using an adaptive threshold value determined by combining both 2D and 3D threshold values as described in (*29*). Chromosome masses were detected after removing very small and scattered pixels by connected component analysis. Maternal and paternal chromosome masses were represented by their three orthogonal Eigen vectors and associated eigenvalues where the eigenvector with the smallest eigenvalue represented the shortest elongated axis of the chromosomal volume. This smallest Eigen value was used as a measure for chromosome congression over time. Since duration of individual phases in different embryos may vary, they were normalised to average phase time. This resulted in uniform phase wise progression among all the embryos and allowed comparison in time. Correlation (Pearson) of the smallest eigenvalues between maternal and paternal chromosomes was computed over time considering a sliding window of size 15.

#### Analysis of directionality of growing microtubule tips

To visualize MT organization we filtered the EGFP-EB3 movies with a Laplacian of Gaussian filter of size 2 px (ImageJ plugin LoG3D, (*30*)). Kymographs were computed for a 11 px wide line using the kymograph analyze feature of ImageJ. EB3 comets have been tracked using the Matlab tool u-track2.0 (Danuser Lab; www.utsouthwestern.edu/labs/danuser/software/, (*31*)). We used the original unfiltered fluorescence images and following parameters: Low pass filter 1 px, High-pass filter 3 px, minimum track length 3, maximum gap to close 1. Due to the 4D nature of the problem the track length was typically short with approximately 80% of the tracks shorter than 5 frames. For the analysis we used tracks longer than 3 frames that are sufficient to determine the overall direction of microtubule growth. This gave over 7000 tracks per embryo. As the tracks were straight we used the vector determined by the initial and end point of the track for further processing. To cluster similar trajectories we performed spectral clustering using the absolute cosine distance between trajectories to construct the similarity matrix (*32*). Then we computed eigenvalues and eigenvectors of the random-walk normalized graph-Laplacians (*33*). The eigengap heuristic indicated that three clusters were sufficient to describe our data (not shown). Finally, to remove isolated tracks we performed a density-based clustering using the average track position (*34*). The *eps* parameter for dbscan was set so to keep 70-80% of the tracks. This gave *eps*-values between 4 and 7 px.

#### Correlation between 5mC and 5hmC signals

Segmentation of 5mC and 5hmC signals was performed using a script developed in MATLAB that quantifies the distribution of the signals and their correlation using bright pixels. Segmentation of maternal and paternal centromeres was performed using an in house developed MATLAB segmentation pipeline and the mixing between parental centromeres was measured using the overlap between their 3D convex hulls.

The Yoyo channel was interpolated first to obtain isotropic voxels. A 3D Gaussian filter of size 3 and SD=2 was applied to reduce the effect of noise. Interpolated stacks were binarized by combining 2D and 3D threshold values and chromosome masses were detected by connected component analysis. Interpolated slices were removed to keep only original slices in order to detect 5mC and 5hmC signals inside the chromosome mass. Spatial correlation between these two signals was performed considering only their bright pixels (top 25%) detected from their histograms. Then their correlation was computed as below

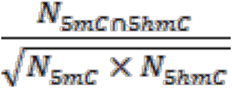

Where *N_smC_* and *N_shmC_* represent the number of 5mc and 5hmC top 25% positive pixels, respectively.

#### Volume occupied by Top 50% of pixels for 5mC, 5hmC, Ring 1B and H3K9me3

Segmentation of 5mC, 5hmC, Ring1B and H3K9me3 regions was performed as described above, however a number of top pixels that added up 50% of the total intensity of the signal were selected. The volume occupied by these pixels was divided by the total volume of the signal.

Average intensity for maternal and paternal pro-nuclei 5mC and 5hmC labelling Chromosome regions were segmented from the DNA channel. The average background intensity was calculated from the intensity in chromosome free regions and was subtracted from the original signal. The average intensity of pro-nuclei was calculated from the background subtracted signal.

---

**Figure S1.**
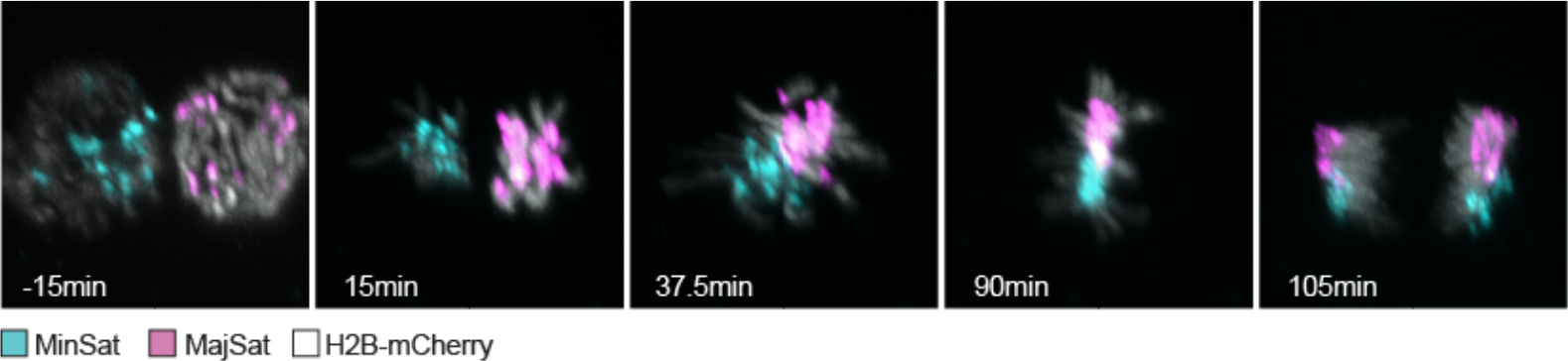
Distribution of parental chromosomes in mouse zygotes. Differential labelling of maternal (magenta) and paternal (cyan) centromeres through distinction of minor and major satellite regions by fluorescent TALEs (*n* = 19). Chromosome arms are labelled with H2B-mCherry (grey). Shown are 3D-rotated images of parental chromosome distribution at different time points during the first mitosis of live imaged MMU x MSP zygotes are shown. NEBD is at time point 0.

**Figure S2.**
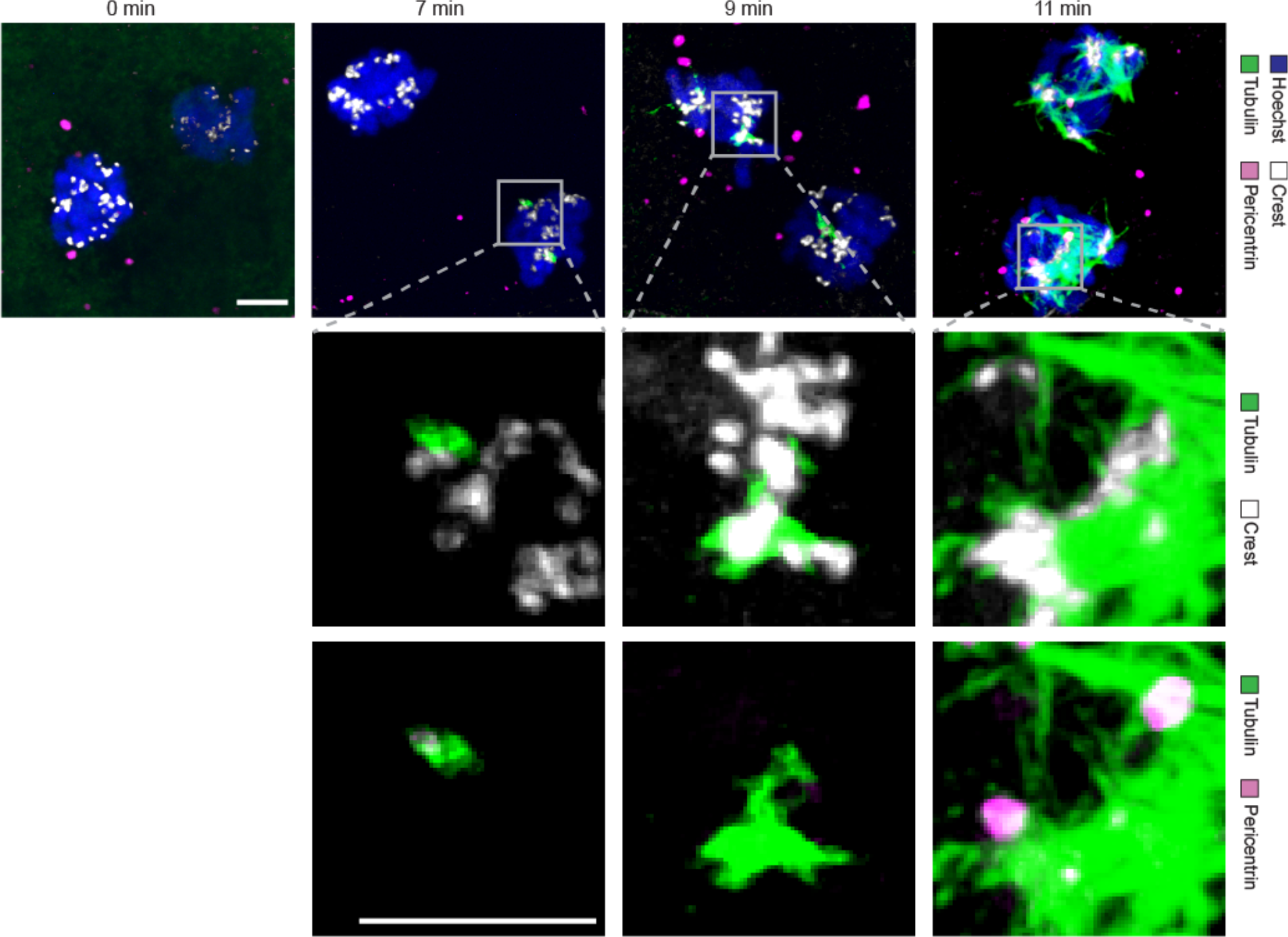
Microtubules are nucleated by the kinetochores in early pro-metaphase. Immunofluorescence staining of MMU x MMU zygotes fixed in 1-minute-intervals up to 13 minutes after Nocodazole washout. Shown are z-projected images of confocal sections of time points 0, 7, 9 and 11 min (*n* **=** 4 per time point, data not shown are consistent with the displayed images). Overview images (first row) with microtubule (tubulin; green), MTOC (Pericentrin; magenta), kinetochore (Crest; grey), and DNA (Hoechst; blue) channels overlaid. Grey boxes, position of magnified images in 2^nd^ and 3^rd^ row. Second row, magnifications with microtubules (green) and kinetochores (white) channels overlaid. Third row, same magnifications with microtubules (green) and MTOCs (magenta) channels overlaid. Scale bars, 5 µm.

**Figure S3.**
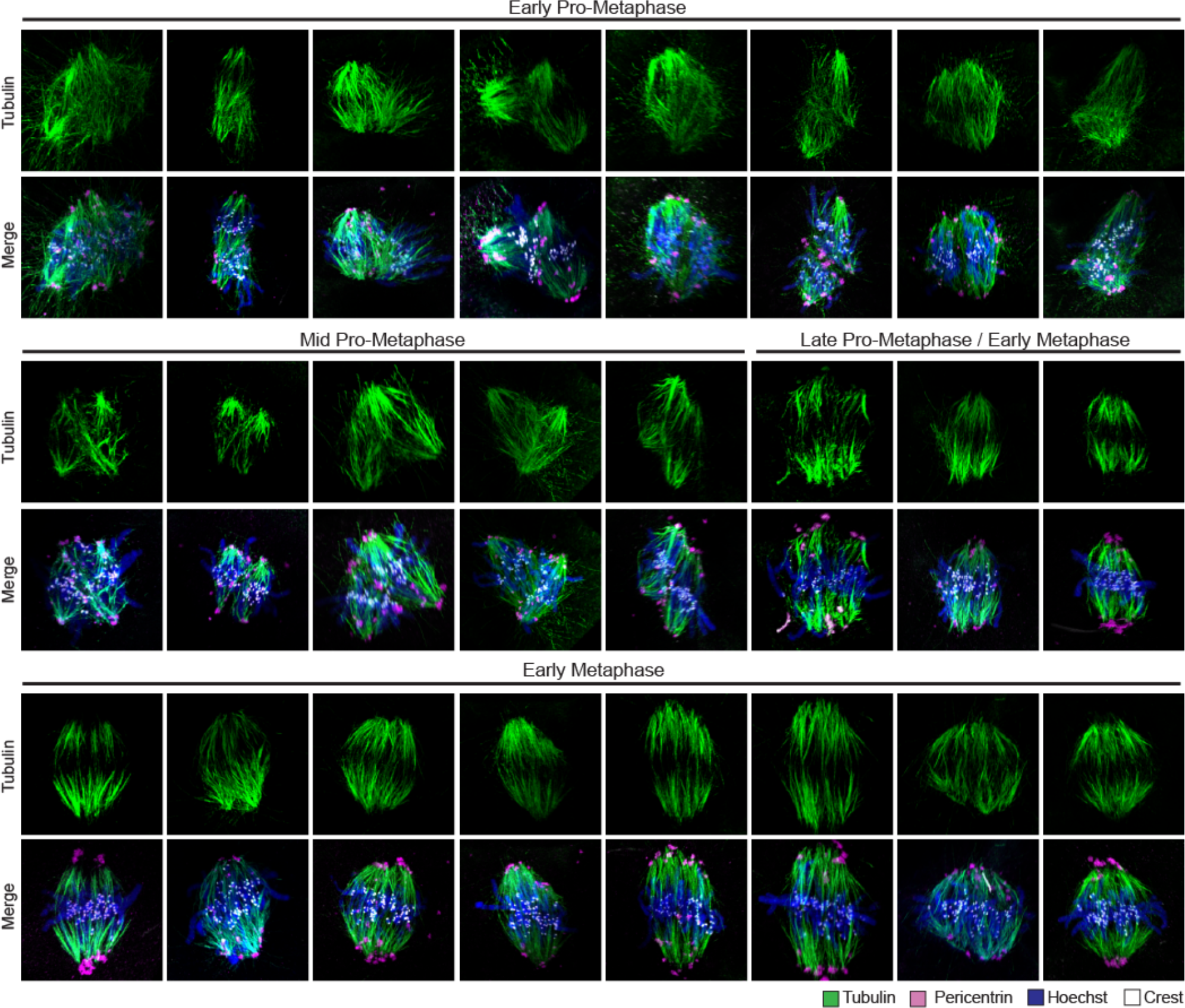
Stages of dual spindle assembly during pro-metaphase. Immunofluorescence staining of cold-treated MMU x MMU zygotes fixed between 10 to 30 minutes after NEBD (*n* = 24). Shown are 3D-rotated images of the whole spindle volume. Embryos are classified as early pro-metaphase, mid-prometaphase and late pro-metaphase/early metaphase based on distance of parental chromosome masses. First rows, microtubules (Tubulin; green) only. Second rows, overlays of microtubule (green), MTOC (Pericentrin; magenta), kinteochore (Crest; grey), and DNA (Hoechst; blue) channels.

**Figure S4.**
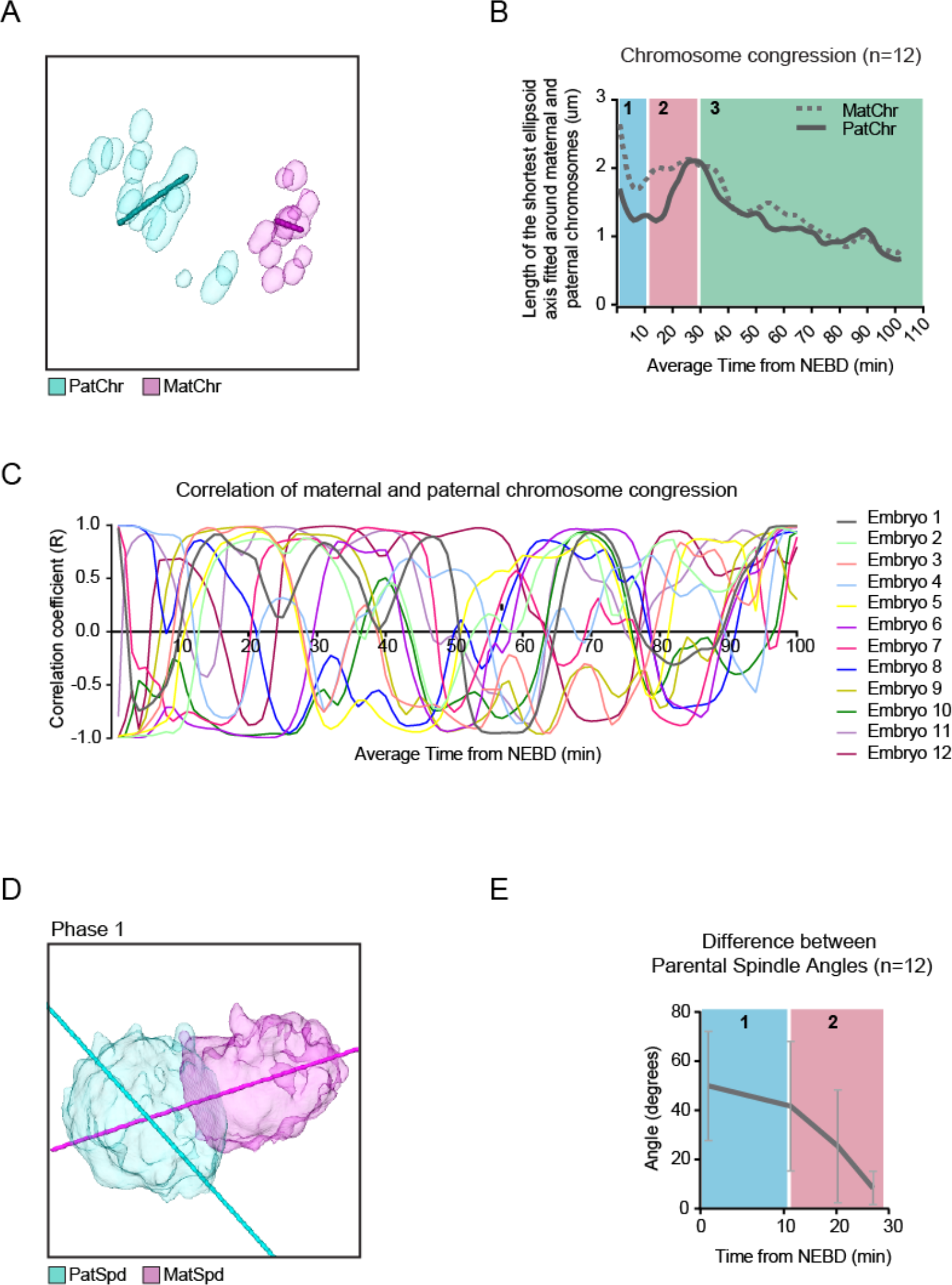
Bipolar spindle formation around each pronucleus. (A) Schematic of ellipsoid fitting to distributions of maternal (MatChr) and paternal chromosomes (PatChr) to determine chromosome congression by measuring the ellipsoid’s shortest axis. (B) Length of the ellipsoids’ shortest axis measurements of maternal (dashed line) and paternal chromosomes (solid line) during phase 1 (blue), 2 (red) and 3 (green). (C) Correlation of maternal and paternal chromosome congression is plotted. Correlation coefficients (*r*) of chromosome congression over time for 12 embryos are shown. (*r* = 0: no correlation, *r* = 1: linear relationship, *r* = −1: negative correlation; for details see methods). (D) Schematic of angle measurements of maternal and paternal spindles. (E) Angles of the long axis of the paternal spindle vs. the long axis of the maternal spindle during phase 1 (blue) and phase 2 (red). (A-E) *n* = 12, MMU x MSP zygotes.

**Figure S5.**
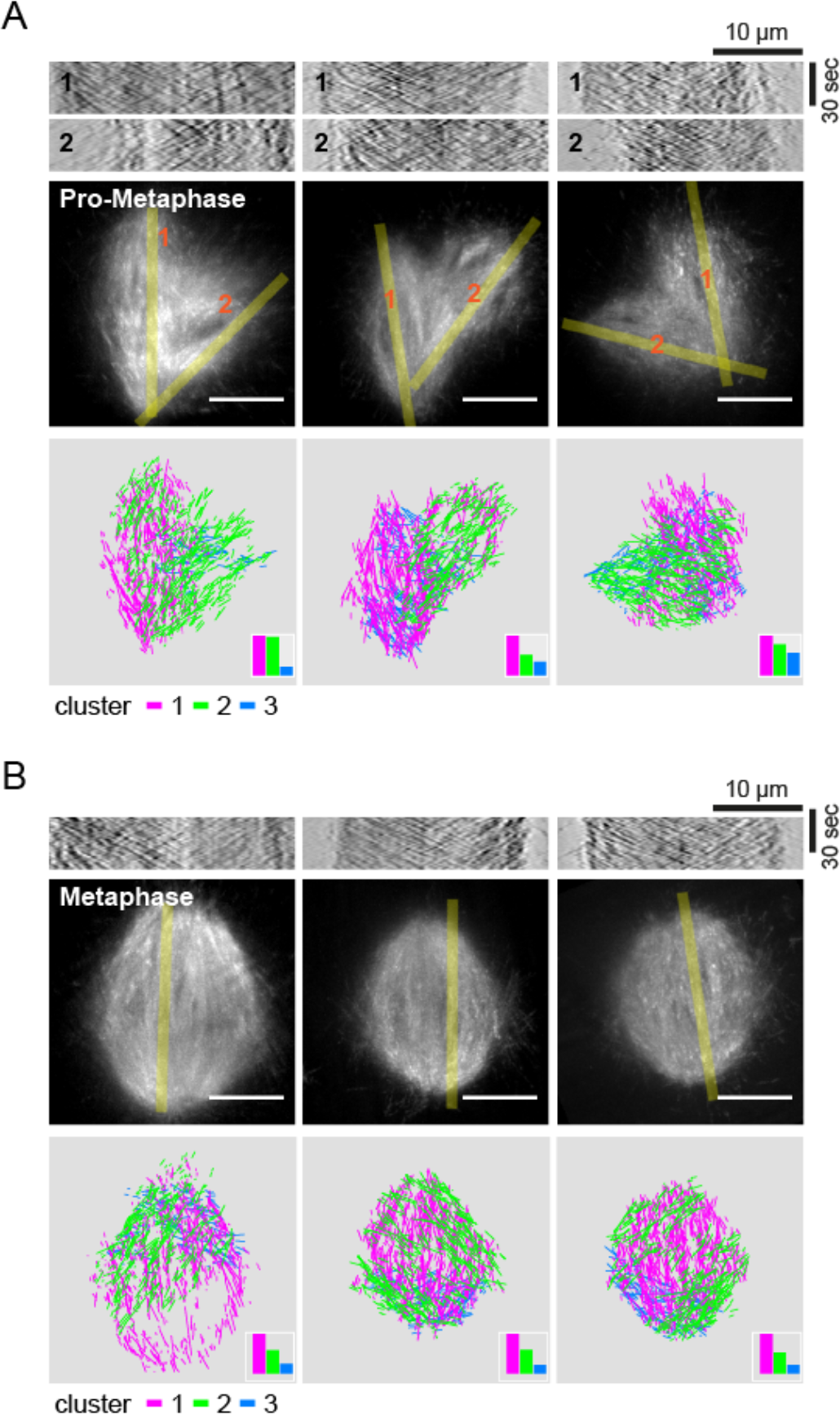
Microtubule (MT) organization in zygotes of MMU x MMU embryos expressing the microtubule tip plus-end marker EGFP-EB3. (A) MT-organization of zygotes in (A) pro-metaphase (phase 2; *n* = 3) and (B) metaphase (phase 3; *n* = 3). (A-B) Upper panels, kymographs of MT flow taken along the yellow lines in middle panels. Middle panels, maximum intensity projection of EGFP-EB3 (white). The yellow line indicates the main axis of the MTs-masses (width 1.43 µm) from which the kymographs are extracted after Laplacian of Gaussian filtering. Lower panels, density-based clustering of MT-tracks (for details see Methods). Three main clusters are shown with the dominant clusters printed in front. For visualization purposes, only 1500 tracks are shown. The insets in the lower panels show the fraction of each cluster. Scale bars, 10 µm.

**Figure S6.**
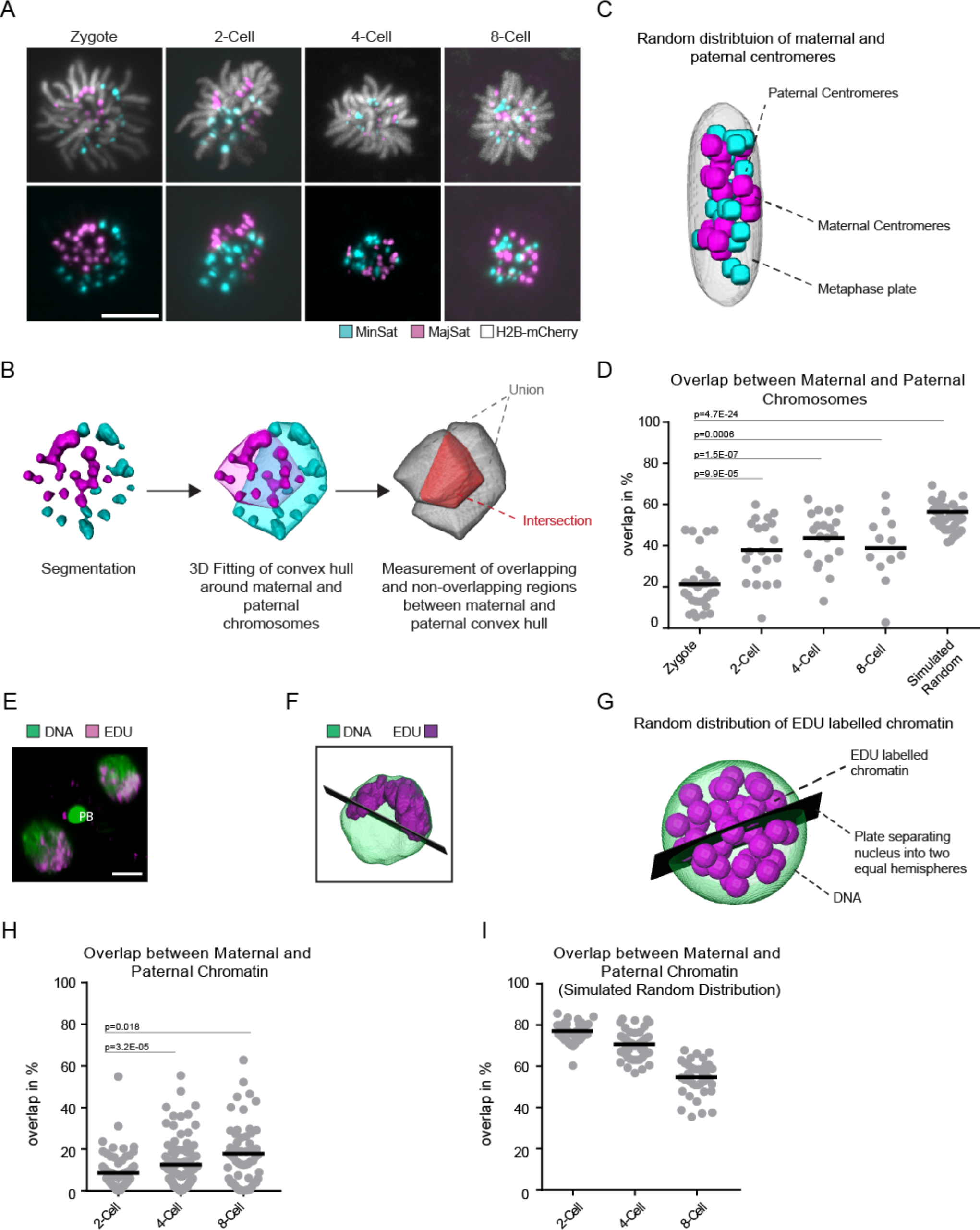
Distribution of parental chromosomes in mouse zygotes. (A) Differential labelling of maternal and paternal centromeres by fluorescent TALEs through distinction of major (MajSat; magenta) and minor satellite regions (MinSat; cyan), repectively. Chromosome arms are labelled with H2B-mCherry (white; in upper row only). Representative *z*-projected images of parental chromosome distribution on the metaphase plate of live-imaged MMU x MSP zygotes, 2-cell, 4-cell and 8-cell embryos are shown. Scale bar, 10 µm. (B) Schematic of 3D convex hull measurement used to determine degree of mixing between parental chromosomes. (C) Schematic of randomized parental centromere distributions at metaphase. (D) Degree of overlap between parental chromosomes at metaphase determined by 3D convex hull measurement for zygotes (n=31), 2-cell (n=20), 4-cell (n=20), 8-cell (n=12) embryos and embryos with in silico randomised distribution (n=40). (E) Distribution of paternal chromatin in MMU x MMU interphase nuclei. EdU-treated male mice were mated with untreated females and the resulting embryos stained for the incorporated EdU with Alexa Fluor 647 (green) by click chemistry and the nucleus counterstained with Sytox green (blue). Representative *z*-projected image for a 2-cell embryo is shown. Scale bar, 10 µm. (F) Schematic of paternal chromatin distribution measurement. (G) Schematic of randomized paternal chromatin distribution at interphase. (H) Degree of paternal chromatin compartmentalization at interphase measured for 2-cell (*n* = 69), 4-cell (*n* = 97), 8-cell (*n* = 55) nuclei. (I) Degree of paternal chromatin compartmentalization at interphase in randomized 2-cell, 4-cell and 8-cell nuclei (each *n* = 40). A-C, E-G, for details see methods. Difference in simulated values between D and H are caused by the different constraints of the arrangement on a metaphase plate or interphase nucleus (sphere) on the distribution of parental signal. (D, H) Statistics, student’s t-test.

**Figure S7.**
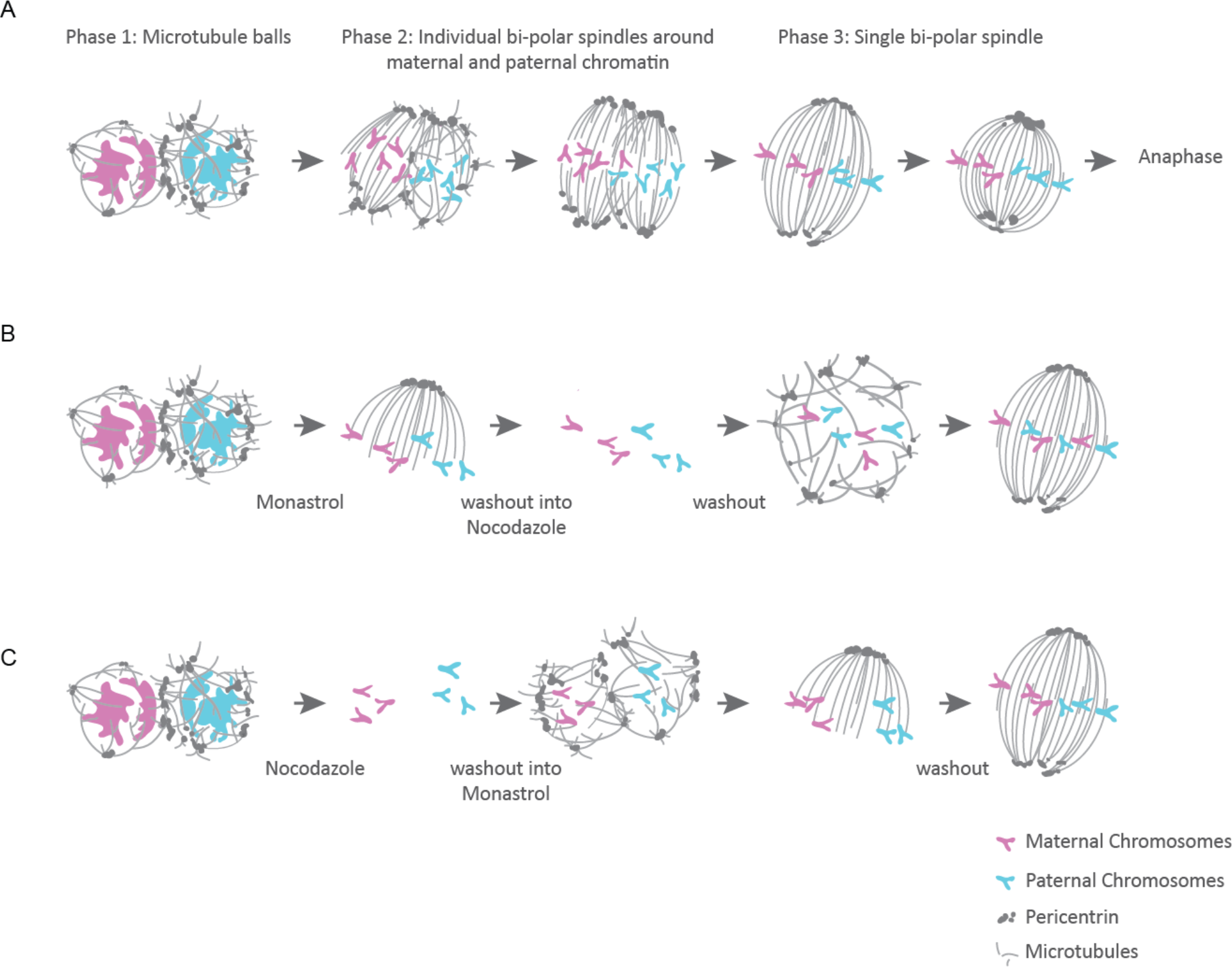
Schematic summary of zygotic spindle assembly and experimental outline for achieving mixing of parental chromosomes during the first embryonic division. (A) Progressive transition from two bipolar spindles to a single barrel-shaped spindle. (B) Treatment of zygotes with Monastrol prior to NEBD brings maternal and paternal chromosomes into close proximity on a monopolar spindle. Subsequent washout into Nocodazole results in depolymerization of microtubules. Washout of Nocodazole allows the spindle to reform in a random orientation leading to mixing of parental chromosomes. (C) Treatment of zygotes with Nocodazole prior to NEBD will result in maternal and paternal chromosomes condensing away from each other. Washout into Monastrol will result in maternal and paternal chromosomes aligning on a monopolar spindle. Washout of Monastrol will cause monopolar spindle to grow into bipolar spindle but separation of parental chromosomes will be maintained.

**Figure S8.**
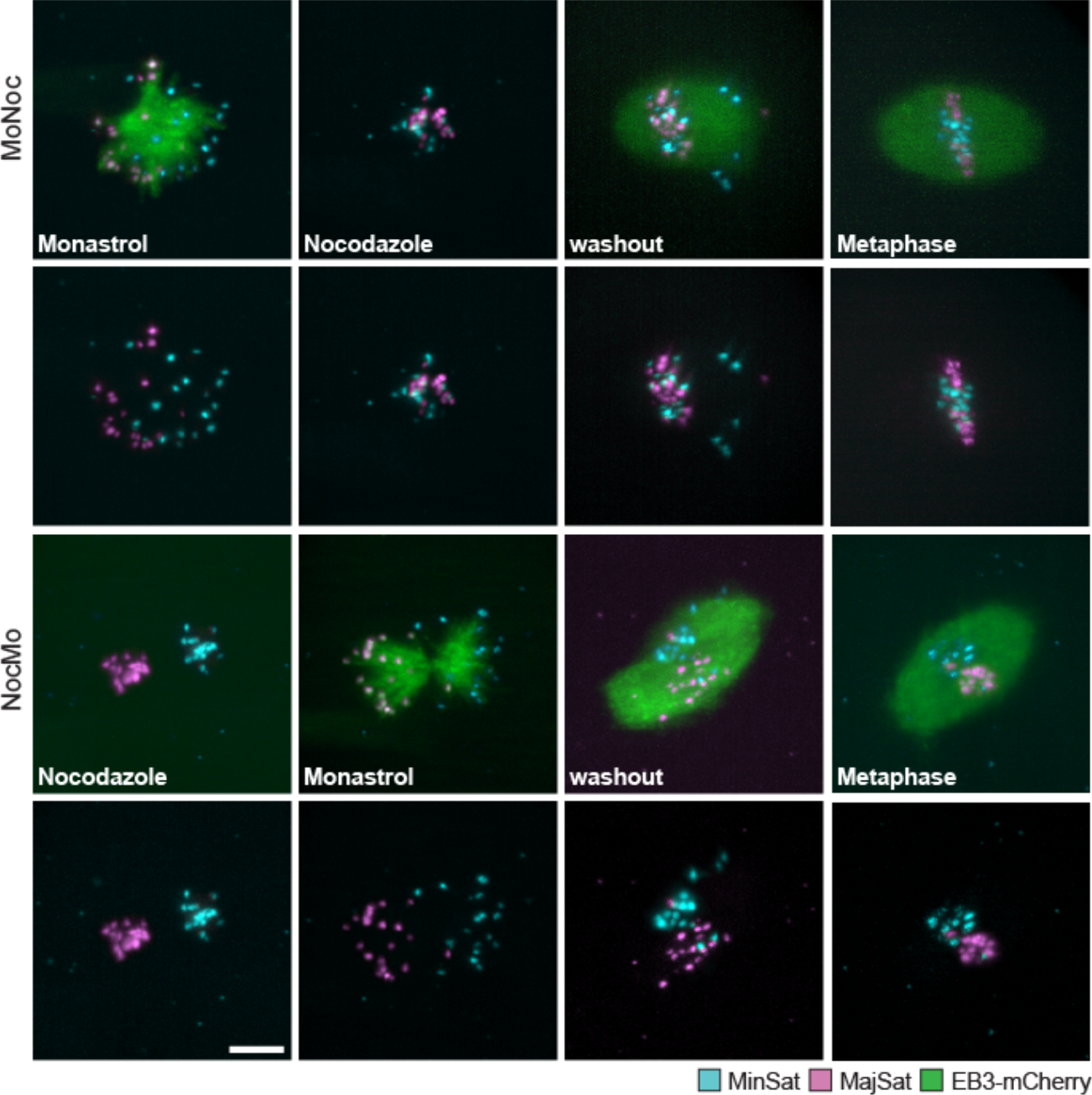
Zygotic spindle formation in NocMo and MoNoc treated embryos. Live imaging of MMU x MSP zygotes expressing fluorescent TALEs to label maternal (MajSat; magenta) and paternal (MinSat; cyan) chromosomes and microtubules (EB3-mCherry; green). The upper two panels show that a single bipolar spindle can be formed when a monopolar spindle collecting both parental genomes is first induced with Monastrol treatment prior to nuclear envelope breakdown, followed by transient microtubule depolymerization with Nocodazole (*n* = 3). The lower two panels show that zygotes in which the drug treatment is reversed form two monopolar spindles upon Monastrol treatment mature into one bipolar spindle upon washout of the drug (*n* = 3). Scale bars, 10µm.

**Figure S9.**
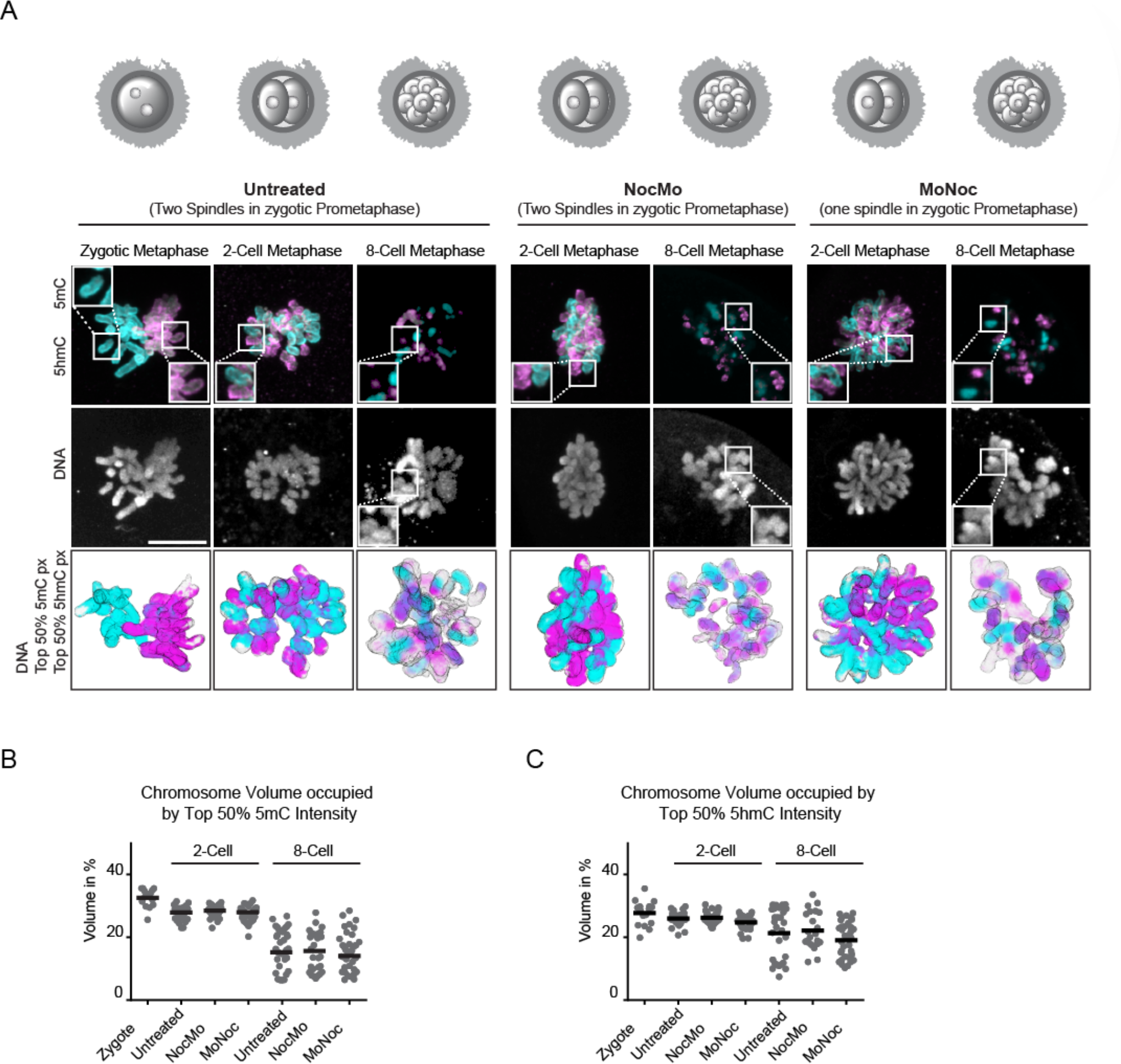
Intensity and distribution of 5mC and 5hmC in mixed and unmixed MMU x MMU embryos. (A) Immunofluorescence staining of zygotes, untreated, NocMo and MoNoc embryos at the 2-cell and 8-cell stage. Shown are *z*-projected images of confocal sections. Top panel shows 5mC and 5hmC staining (magenta, cyan) (salt and pepper distribution is expected based on results on parental chromosome distributions as determined in Fig. S1 at the 2-cell stage). 5mC and 5hmC intensities were individually scaled per image to show top 50% of 5mC and 5hmC positive voxels, respectively. Middle panel shows DNA staining (white). Bottom panel shows chromosome surface (grey) and top 50% intensity pixels of 5mC (magenta) and 5hmC (cyan) printed as solid colors (see methods for details). Scale bar, 10 µm. (B) Volume occupied by top 50% 5mC positive voxels. (C) Volume occupied by top 50% 5hmC positive voxels. Zygote (n=16), untreated (n=26), NocMo (n=47) and MoNoc (n=39) cells at the 2-cell stage and untreated (n=30), NocMo (n=22) and MoNoc (n=35) cells at the 8-cell stage.

**Figure S10.**
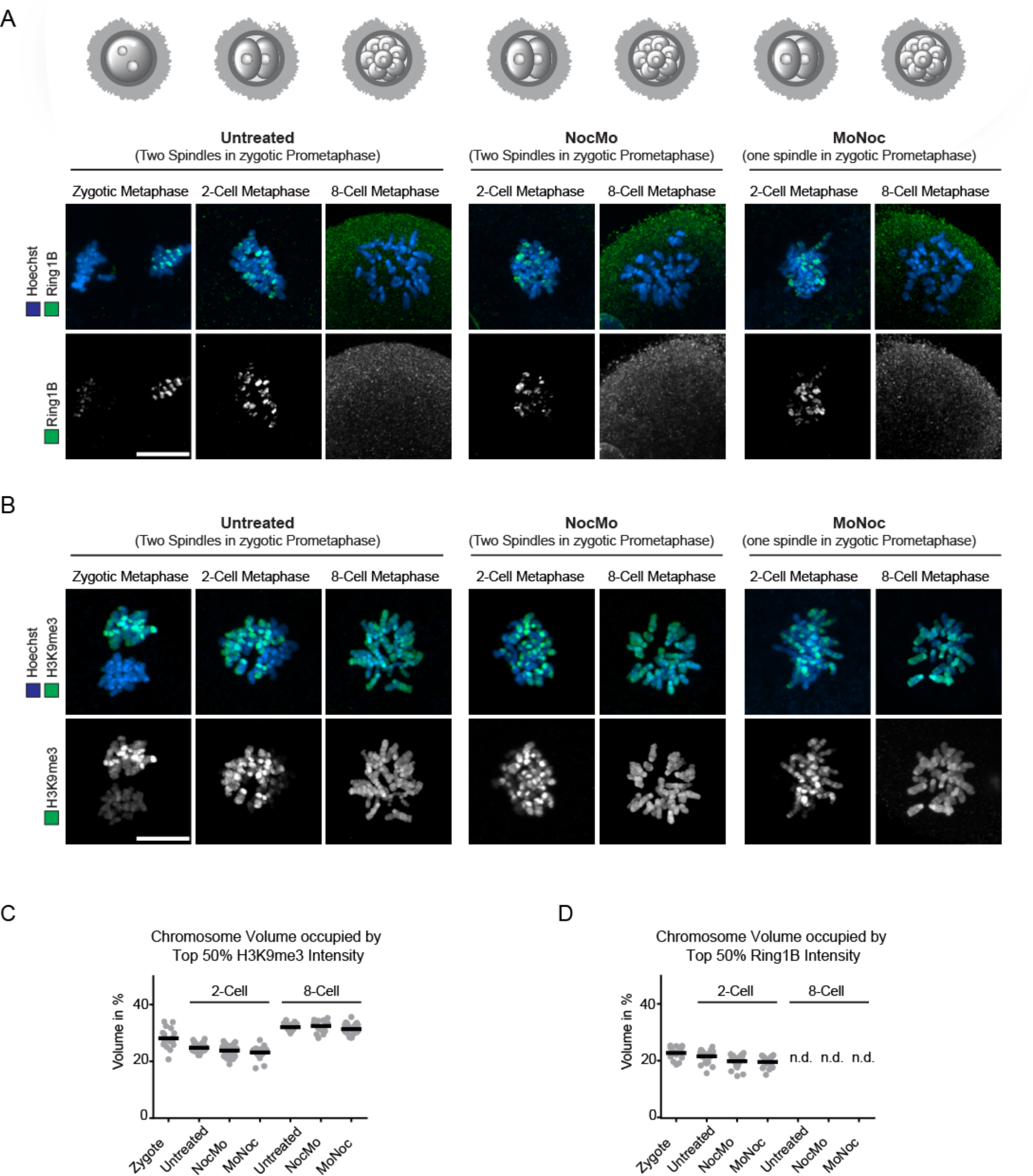
Intensity and distribution of Ring1B and H3K9me3 in mixed and unmixed MMU x MMU embryos. (A) Immunofluorescence staining of zygotes, untreated, NocMo and MoNoc embryos at the zygote, 2- and 8-cell stage showing DNA (Hoechst; blue) and H3K9me3 (green). Shown are *z*-projected images of confocal sections. H3K9me3 intensity was individually scaled per image to show top 50% of positive voxels. (B) Immunofluorescence staining of zygotes, untreated, NocMo and MoNoc embryos showing DNA (blue) and Ring1B (green). Shown are *z*-projected images of confocal sections. Ring1B intensity was individually scaled per image to show top 50% of positive voxels. (C) Volume occupied by top 50% H3K9me3 positive voxels. Zygote (*n* = 20), untreated (*n* = 31), NocMo (*n* = 41) and MoNoc (*n* = 22) cells at the 2-cell and untreated (*n* = 33), NocMo (*n* = 44) and MoNoc (*n* = 33) cells at the 8-cell stage. (D) Volume occupied by 50% Ring1B positive voxels. Zygote (*n* = 17), untreated (*n* = 37), NocMo (*n* = 34) and MoNoc (*n* = 19) cells at the 2-cell. For 8-cell stage, volume of positive voxels was not determined due to absence of signal. Scale bar, 10 µm. Furthermore, we re-implanted control, MoNoc and NocMo embryos into foster mothers and show that similar numbers of pups get born from each condition (Table S1).

**Table S1.**
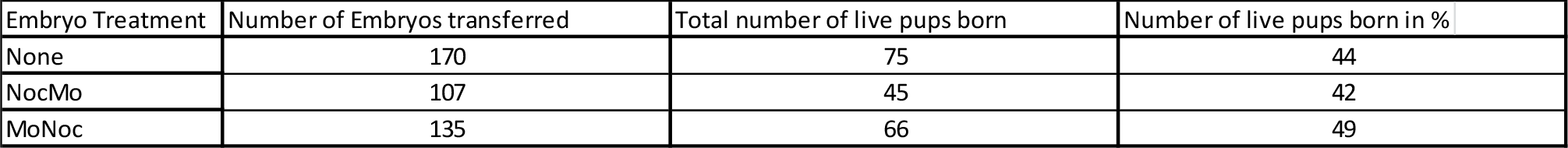
Frequency of pups born after re-implantion for untreated zygotes, zygotes treated transiently with Nocodazole followed by Monastrol (NocMo) and zygotes treated transiently with Monastrol followed by Nocodazole (all zygotes were collected from MMU x MMU crosses).

**Movie S1.** Live-cell time-lapse imaging of MMUxMSP mouse zygote expressing fluorescent TALEs for differential labelling of maternal (magenta) and paternal (cyan) centromeres through distinction of minor and major satellite regions. Chromosome arms are labelled with H2B-mCherry (grey). Time resolution is 7.5 min.

**Movie S2.** Live-cell time-lapse imaging of MMU mouse zygote expressing EB3-mCherry (green) and tdEos-Cep192 (magenta). Time in min.

**Movies S3-5.**

Live-cell time-lapse imaging of pro-metaphase mouse zygote (phase 2) expressing EGFP-EB3 Left panel: Maximum intensity Z projection. Right panel: The movie has been filtered with a Laplacian of Gaussian (2×2 px) to enhance the EB3 tip signal (right panel). Scale bar 10 µm. Time resolution is 800 ms.

Movies S3-5 represent the three examples corresponding to Figure S5A.

**Movies S6-8.**

Live-cell time-lapse imaging of metaphase mouse zygote (phase 3) expressing EGFPEB3 Left panel: Maximum intensity Z projection. Right panel: The movie has been filtered with a Laplacian of Gaussian (2×2 px) to enhance the EB3 tip signal (right panel). Scale bar 10 µm. Time resolution is 800 ms.

Movies S6-8 represent the three examples corresponding to Figure S5B.

**Movies S9-11.**

Live-cell time-lapse imaging of MMU mouse zygote after washout of Nocodazole (> 10 h treatment) expressing αTubulin-EGFP (green) and H2B-mCherry (magenta). Time in min. Movies S9-11 represent three examples corresponding to Figure 3.

